# White matter reorganization of motor and affective-motivational networks in pain-indifferent carriers of the R221W mutation

**DOI:** 10.1101/2025.07.09.663977

**Authors:** Arnas Tamasauskas, Irene Perini, Jan Minde, Simon S. Keller, Nicholas Fallon, Bernhard Frank, India Morrison, Andrew Marshall

## Abstract

Congenital insensitivity to pain (CIP) due to the R221W mutation on the nerve growth factor gene results in reduced peripheral C-nociceptor density and behavioural indifference to painful stimuli. While functional neuroimaging has revealed altered cortical and sub-cortical pain processing in R221W carriers, structural white matter changes remain unexplored and may suggest an anatomical basis of symptoms. Heterozygous R221W carriers’ (n = 11) and age-, sex-, education-matched controls’ (n = 11) diffusion tensor imaging data were compared using fixel-based analysis, and complimentary edge and node analyses using graph theory, and network-based statistics. Whole-brain and region of interest (ROI) fixel-based analyses revealed significantly reduced fibre density and fibre-bundle cross-section in brainstem motor tracts of R221W carriers, encompassing the corticospinal pathways, corona radiata, external capsule, cerebellar peduncles, and pontine crossing (p < 0.05). Graph theory analysis of pain-processing ROIs demonstrated reduced local efficiency in right anterior cingulate cortex (ACC) and altered betweenness centrality in bilateral insula and left ACC of R221W carriers. Despite R221W carriers showing higher node degrees in the somatosensory cortex and ACC, these connections had reduced efficiency and integration with cortical network regions. Network-based statistics identified a possible compensatory subnetwork with stronger connectivity from right thalamus to left ACC and left insula in R221W carriers (p < 0.019). These findings suggest that congenitally reduced peripheral nociception could lead to abnormalities in the thalamocortical and motor efferent pathway, but not sensory afferent pathways. The combination of reduced brainstem motor tract integrity and altered cortical network efficiency, alongside potentially compensatory thalamo-cortical connectivity, could support a model of R221W CIP as motor under-reactivity rather than sensory insensitivity.

## 1. Introduction

For most of us, pain is an unescapable part of being. However, for rare individuals with reduced pain sensitivity, pain is just a subtle, insignificant sensation. Acquired peripheral neuropathies usually result in slightly reduced pain sensitivity^1,2^, such as reduced reactivity to pinpricks. Congenital Insensitivity to Pain (CIP)^3,4^ syndromes are more severe, resulting in individuals sustaining significant bodily harm and letting their wounds go untreated, particularly broken bones and fractures^3^. The prevalence of CIP is unknown; some studies have reported the prevalence of 1 in 25,000^4^ but the origin of these figures in scientific literature is unclear.

CIP has been seen in people with mutations in SCN9a^5^, CLTCL1^6^, NTRKI^7^, and R221W^8^ genes. The R221W gene mutation in particular results in reduced nerve growth factor neuropeptide secretion. This reduction results in a lower density of unmyelinated C-nociceptor afferent nerves in the skin – a type of peripheral neuropathy^,9^. C afferents carry somatosensory information of mechanical, thermal, and chemical nociception from the body’s periphery to the central nervous system^10^. Recent research into R221W carriers has shown that this reduction in unmyelinated C-afferents results in indifference to pain^11,12^. These afferents form part of nociceptive pathways and the reduced activity translates to under-activation of pain-related brain areas, possibly linked to carriers’ observed behavioural under-reaction to pain. While R221W heterozygote carriers are able to report subjective thermal pain thresholds to the same degree as non-carrier controls, they lower and slower reported motivational urge to escape painful heat stimulation compared to controls^11^. These behavioural responses correlated to an under-activation of the middle part of Anterior Cingulate Cortex (ACC) and anterior insula during a functional Magnetic Resonance Imaging (MRI) motor response task involving heat pain stimulation^13^. These findings imply that altered peripheral nerve density is related to altered pain processing in the heterozygote R221W carrier population. As research has shown that ACC and anterior insula areas are usually linked to a participant’s motivation to respond; it was hypothesized that R221W carriers showed increased reliance on the sensorimotor cortex instead^13^.

Functional MRI activation changes were also seen in people with acquired peripheral neuropathy. In diabetic neuropathy, patients can present with hyperalgesia or hypoalgesia sensory signs^14,15^. When these groups were compared in a heat pain functional MRI task, people with hypoalgesia exhibited reduced functional connectivity between the insula, amygdala and ACC^16^. This reduced functional connectivity correlates with white matter connectivity changes observed in people with diabetic neuropathy. Using structural MRI, people with diabetic neuropathy were shown to have reduced white matter pathways in the sub-cortical pain processing areas including the thalamus, insula, and ACC compared to healthy controls.^17,18^ In cases of peripheral neuropathy, it is difficult to discern whether these structural and functional changes in the brain are due to under-activated peripheral-central nerves, or whether these changes are due to neurological damage from the conditions that have caused neuropathy^19^. For example, acquired neuropathy due to diabetes is commonly associated with small vessel disease or chronic inflammation^19,20,21,22^. Cases of CIP reduce these health cofounders and provide a unique opportunity to understand the direct impact of peripheral neuropathy on the development of abnormalities within central nervous system.

To understand central nervous system alterations in R221W carriers, diffusion tensor imaging (DTI) can provide insight into structural connectivity anomalies^23^. However, DTI modelling has historically been limited by statistical, practical and theoretical uncertainty in how to approach crossing fibres^24^. Fixel-based analysis provides a novel technique that quantifies fibre orientation distribution in individual voxels into “fixels”^25,26^ (a fibre equivalent of voxels), which represent individual fibre populations within a given voxel, allowing for increased specificity over voxel-wise measures. Using fixel-based analysis, precise fibre bundles within crossing-fibre voxels can be compared using group-wise analyses of three metrics: fibre density (FD), fibre-bundle cross-section, and a combined measure of fibre density and fibre-bundle cross-section (FDC),^27^which may be reflect different aspects of neural damage. Fixel-based analyses have been shown to be more sensitive to pathological alterations of white matter in various neurological disorders compared to analysis of conventional DTI scalar metrics ^28,29,30^. These analyses often include both whole-brain analysis that provides a more exploratory approach, and a deterministic region of interest (ROI) analysis that provides a more targeted approach. Building on functional MRI research of R221W carriers^13^, the ROI for this study was defined to include key cortical pain processing regions: left and right sensory cortices, left and right ACC, left and right insula, left and right thalamus, and brainstem^31,32,33,34,35,36^. As this was a first investigation of white matter tracts in a CIP cohort, the exploratory whole-brain analysis provided supplementary exploration of areas that are not directly related to pain processing.

Alongside fixel-analysis, ROI areas were analysed using graph theory – a computational neuroscience analysis that provides information of how well vertices of converging connections, referred to as ‘nodes’, are integrated, segregated, or centralised in brain networks^37,38,39,40^. In graph theory, nodes are vertices of converging connections linked by edges which represent connectivity of nodes in a network, allowing analysis of how a brain network is organised. Graph theory provides a complimentary approach to fixel-analysis, as it can be used to investigate how focal changes, that are identified by fixel-analyses, relate to topological differences in pain processing networks. These topological differences reflect the magnitude and importance of nodes and edges connecting them. Edges can additionally be analysed on a group level using Network Based Statistics (NBS)^41^. This method utilises a nonparametric permutation approach combined with cluster-based thresholding to identify suprathreshold links. NBS has been instrumental in identifying edge connectivity differences, that were not previously found in graph theory research, in peripheral neuropathy groups. Graph theory has identified a potential link between acquired painful diabetic neuropathy and increased global and local efficiency in the ACC and reduced in the insula^42,43,44^. Research using NBS on a painful diabetic neuropathy group found an almost opposite effect – the insula had reduced connectivity to cortical pain processing areas^45,46^. The contrasting NBS and graph theory findings on diabetic neuropathy participants’ insula connectivity illustrates how both methods identify complimentary but distinct topological changes that provide nuanced connectivity findings. While these investigations provide information about the structural reorganisation in chronic pain groups^45,46^, there is a no investigation of the opposite sensation – neuropathy with hypoalgesia. Graph theory and NBS together can provide a robust investigation into R221W carriers’ global brain network reorganisation, and provide wider white matter connectome context to fixel-based analysis.

The aim of the present study was to identify how structural brain networks correlate to congenitally reduced C-nociceptive fibres. Fixel-based whole-brain and ROI analyses were carried out to investigate CIP individuals’ brain structural white matter integrity and provide structural context for previous functional MRI findings. Graph theory and NBS analyses can shed light on how focal changes correlate to network organisation in shaping somatosensory signalling and perceptual experience. It can also pave the way for new approaches for interrogating this role, and of further testing and interpreting effects of CIP on central nervous system phenotypes. As there is no previous DTI research into congenital pain insensitivity, the hypotheses of this study were based on functional MRI R221W research^12^ and DTI research of another peripheral neuropathy group – diabetic neuropathy^47^. We hypothesised that R221W carriers would show reduced fibre density and cross section in the corticospinal and spinothalamic tract when compared to healthy controls. We hypothesised that ROI analysis would show reduced integrity of R221W carriers’ supplementary pain processing white matter tracts – cingulum, and anterior limb of internal capsule. We additionally hypothesised that in R221W carriers, graph theory and NBS would show integration, segregation, and centrality to be increased in ACC and Insula but reduced in sensory cortices.

## 2. Methods

### 2.1 Participants

The study protocol was in accordance with Declaration of Helsinki and approved by Gothenburg University Hospital ethics board as a collaboration with Umeå Univeristy. A group of twelve heterozygous R221W carriers and twelve control participants were recruited for this study. R221W carriers were recruited based on pedigree (at least one carrier parent), and the presence of the mutation ((an arginine-to-tryptophan substitution on one allele of the beta subunit of the NFG gene) later confirmed by DNA sequencing. Arthropathies in R221W carriers show a progressive trend, with mild or unaffected carriers showing clinical symptoms decades later. Both groups were recruited in Sweden and the carriers lived in a geographically dispersed area of northern Sweden.

The R221W carriers group and control group were age, sex, and education matched. The mean age of R221W carriers was 36.2 years, and the mean age of matched controls was 35.8 years. Both groups had 7 female and 4 male participants. Both carriers and controls had 12 years of formal education. Some of the heterozygous carriers exhibited subclinical signs of arthropathies, carpal tunnel, and painless fractures. Cognitive functions and basic reflexes were normal in carriers, and they did not differ from controls on the ability to discriminate noxious and innocuous thermal stimulation as assessed by threshold testing^13^.

### 2.2 MRI Acquisition

Data was acquired using 3T General Electric scanner with 32-channel head coil at the Umeå Centre for Functional Brain Imaging, Umeå University Hospital (Umeå, Sweden). Diffusion acquisition consisted of 32 directions at *b* = 1,000 s/mm–2 and six *b* = 0 s/mm^−2^ volumes acquired at 256 x 256 x 64 mm field of view (TR = 8 s, TE = 0.0814 s). T1-weighted structural images were acquired with the following parameters: voxel size 0.4883 x 0.4883 x 1 mm (TR = 0.0082 s, TE = 0.0015 s), flip angle = 12°, slices = 64, slice thickness=1 mm, field of view= 256 × 256.

### 2.3 Pre-processing

As DTI was acquired using single phase-encoding (anterior-posterior), a generative Synb0-DISCO^48^ algorithm was used to synthesise a reverse (posterior-anterior) phase-encoded image. The acquired DTI and synthesised DTI images were then used for preprocessing using FMRIB software library (FSL)^49,50^. FSL was first used with the TOPUP command to estimate and correct susceptibility induced distortions, and then EDDY command to correct for eddy currents and head movement. MRtrix^51^ toolbox (*dwibiascorrect*) was used for bias field connection and global intensity normalisation across subjects.

The T1-weighted image of each participant was pre-processed using MRTrix^51^ to create a T1-weighted image with five different tissue types – cortical grey matter, subcortical grey matter, white matter, cerebrospinal fluid, and pathological tissue. For each participant, the segmented T1-weighted image was registered to their pre-processed DTI image.

### 2.4 Computation of Fixel Metrics

Recommended methodology was followed for Fixel metrics generation^52^. Pre-processed DTI were bias field corrected using Advanced Normalization Tools^53^. The following steps were then executed using MRtrix^51^ toolbox: DTI images were unsampled and Fiber Orientation Distribution (FOD) estimations was performed with a multi-tissue constrained spherical deconvolution algorithm (*SS3T-*CSD)^54^ using the group average white matter response function. These FODs for each tissue were bias field corrected and had global intensity normalisation (*mtnormalise*) applied across subjects. An FOD template was created from all subjects, and wraps were computed to register each participant to the templates. A Fixel mask was then created from the FOD template. Each individual’s segmented T1-weighted images were also warp registered to the FOD template, and were then used to create an anatomically constrained mask.

For each subject, FOD images were segmented to identify the number and orientation of Fixels in each voxel, as well as estimating apparent FD in each Fixel. Once the directions of all subject image Fixels were reoriented based on a Jacobian matrix, the FD value of each subject’s Fixels were assigned to the corresponding Fixel in the FOD population template. Fibre cross-section was computed from warps generated in previous steps, which was then used to compute a combined FDC metric. Log metric was calculated for fibre cross-section scores to ensure normal distribution (LogFC).

### 2.5 Whole-Brain Fixel-Based Analysis

Whole-brain fibre tractography was performed using a probabilistic algorithm - Second-order Integration over Fiber Orientation Distributions (iFOD2)^55^ with an anatomically constrained mask. Ten million streamlines were generated using the recommended tracking angle (22.5°) and lengths (min 10; max 250). This was followed by Spherical-deconvolution Informed Filtering of Tracks (SIFT)^56^ to control for track thickness overestimation and straight track density bias. For reduction of streamlines using SIFT, a factor of ten was selected as it is a heuristic guideline outlined in MRTrix^57^ and commonly used in Fixel research^58^. A Fixel-Fixel connectivity matrix was then generated from one million streamlines. Fixel connectivity filtering using the connectivity matrix was applied to FD, LogFC, and FDC to reduce noise^59^. Permutation testing was then performed on these metrics with family-wise error rate correction using the threshold-free cluster enhancement^60^. Resulting significant Fixel plots were converted to voxels and used as masks to reduce the whole-brain tractogram to ten thousand streamlines using SIFT for visualisation of white matter streamlines going through significant Fixel areas. These significant Fixel plot streamlines were then registered to a John Hopkins University (JHU) ICBM-DTI-81 white-matter labels atlas^61^ using FSL Linear Image Registration Tool^62^ and Non-Linear Registration Tool^50^ to identify associated anatomical pathways.

### 2.6 ROI Fixel Analysis

ROI analysis was undertaken alongside whole brain analysis to increase sensitivity for findings in a small sample, and to create Fixel parcellated ROI connectomes to be used in graph theory analysis. ROI analysis was performed in brain areas with high likelihood of engagement by pain stimulation^34^: bilateral primary and secondary somatosensory cortices, bilateral ACC, bilateral insula, bilateral thalamus, and brainstem^31,32,33,34,35,36^. Freesurfer^63^ was used to create parcellation images of each participant’s segmented pre-processed T1-weighted scans. The extracted ROIs were warped to standard space and then combined into a mask for each participant. These masks were then averaged using voxel-wise mean to create a pain pathway mask for ROI focused Fixel-based analysis.

Ten million streamlines passing through the pain pathway ROI mask were generated using iFOD2 for each participant. SIFT was performed with a factor of 10, and the resulting million streamlines were used in structural connectivity (SC) matrix generation for each participant. Additionally, Fixel group comparison analysis was performed by using iFOD2 and SIFT on the population template with the pain pathway ROI mask. Analysis of FD, LogFC, and FDC followed the same steps as whole-brain analysis.

### 2.7 Graph Theory and NBS

The resulting node assignments from SC matrix generation were processed to calculate centroid nodes. SC matrices were processed by applying proportional thresholding to keep the top 20% strongest connections to produce edges for analysis. These centroid nodes and edges were then used to calculate graph theory metrics with Matlab Brain Connectivity Toolbox^64^. Following a published systematic review of graph theory in pain neuroimaging^38^, integration was measured by global efficiency, which reflects an inverse of the shortest path to all areas in the network. Segregation was measured by local efficiency, which reflects how well the area is connected to its neighbours. Centrality was measured by node degree, which compared the number of connections to the area, and betweenness centrality, which compared how many shortest paths connect through the area. Efficient local structural connectivity in human brain networks tend to combine high integration (global efficiency) with high segregation (local efficiency). The connections between nodes are referred to as ‘edges’, and their analysis can provide complimentary information on centrality and strength of neural connection coming from nodes. Edges were measured by edge betweenness. All graph theory analyses were permutation-based false discovery rate (FDR) tested. Additionally, edge strength was analysed using Network-Based Statistics (NBS)^65^. NBS constitutes nonparametric testing that controls for family-wise error rate and uses 5000 permutations with t-test thresholds set at 2.5, 3.0, 3.5, and 4.0. The significant P < 0.05 was corrected for multiple comparisons using NBS correction. The NBS process is illustrated in Figure 1.

**Figure 1:**
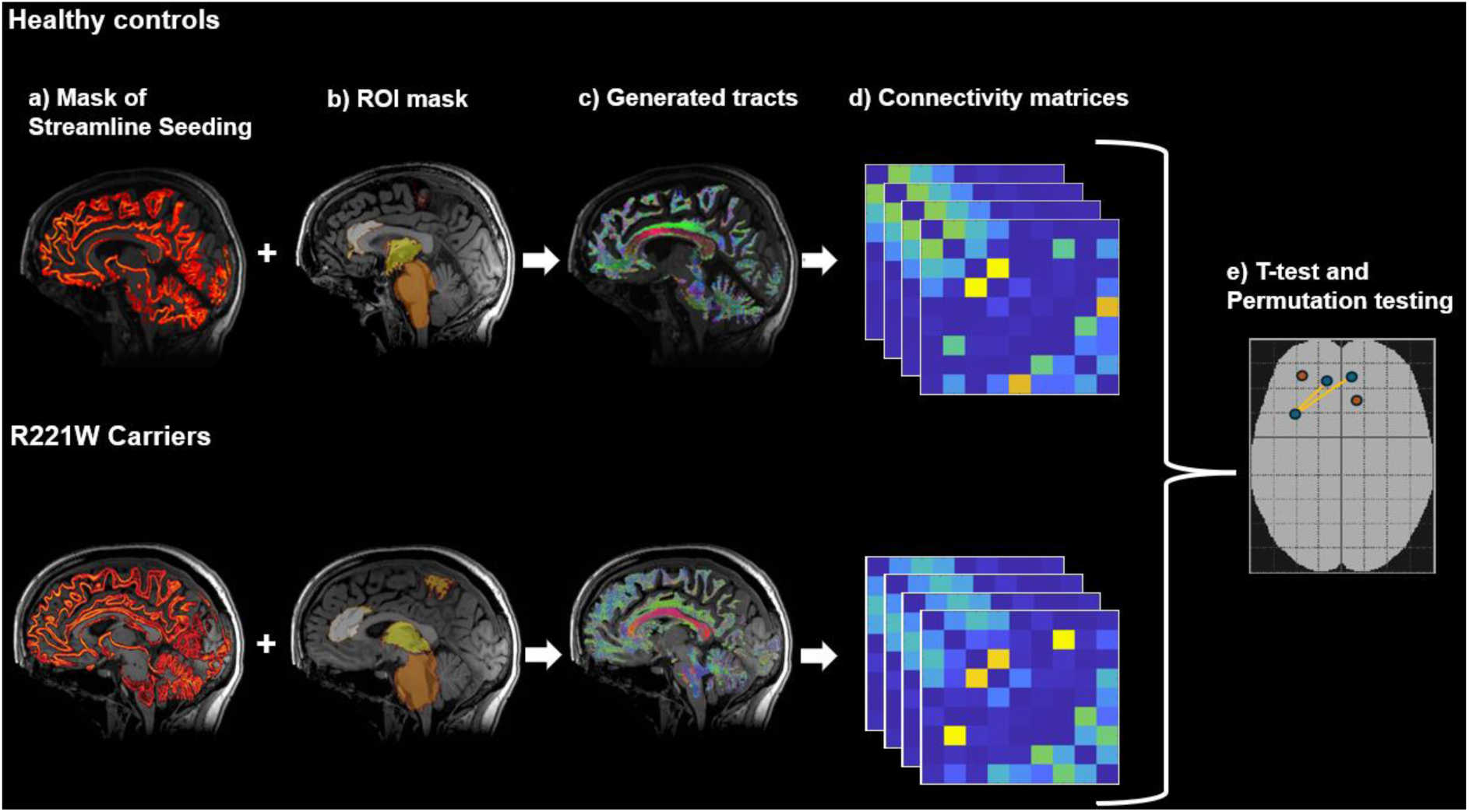
Process of (a) creating anatomically constrained white matter mask with (b) region of interest (ROI) mask to generate (c) Fixel tracts. These tracts then produced (d) connectivity matrices for both groups that were (e) compared using NBS t-test and permutation testing

## 3. Results

### 3.1 Whole-Brain Fixel-based analysis Results

Fixel-based group comparison between R221W carriers and healthy controls showed significantly reduced FD and FDC in midbrain and pons of R221W carriers (p < 0.05) (Figure 2), but no significant differences in LogFC. Overlaying the results on a JHU atlas-based white-matter provided specificity in identifying tracts with significantly reduced FD and FDC of R221W carrier group as compared to healthy controls. These affected tracts were: the middle cerebellar peduncle, corticospinal tract and medial lemniscus (both associated with corticospinal pathways), corona radiata, external capsule, and inferior and superior cerebellar peduncles. FDC revealed only differences in tracts connecting the internal capsule, and uncinate fasciculus. FD and FDC Fixel plots with significance *p* and streamlines, which are coloured according to effect sizes, are visualised in *Figure 2*.

**Figure 2:**
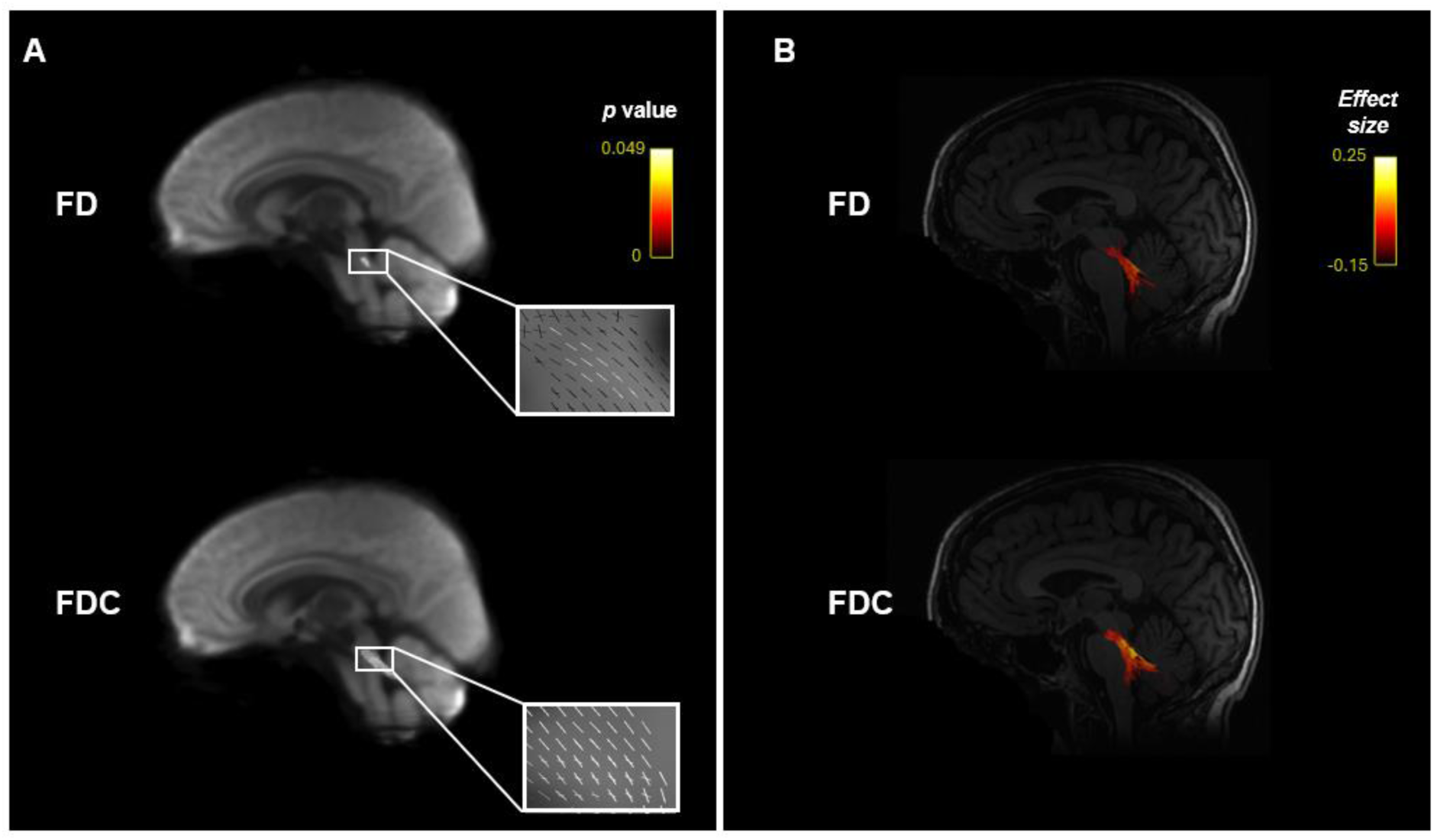
A) Fibre Density (FD) and Fibre Density and Cross-section (FDC) Fixel plots coloured by significance p overlayed over the population white matter template. B) Streamlines coloured by effect side passing through significant Fixel plot FD and FDC areas overlayed over T1 scans.

### 3.2 ROI Fixel-Based Analysis Results

The whole-brain analysis results were replicated in cortical pain-associated ROIs. The Fixel-based group comparison found significantly reduced FD and FDC in the midbrain and pons of R221W carriers (p<0.05). There were no significant LogFC differences. When streamlines passing through significant FD and FDC plots were overlayed over the JHU white matter atlas, most tracts mirrored whole-brain analysis - the corticospinal tract, medial lemniscus, inferior and superior cerebellar peduncles. Additionally, ROI analysis identified significant FD and FDC reduction in the pontine crossing tract. FD and FDC Fixel plots, as well as streamlines going through them, are visualised in *Figure 3* with *p* significance and effect size.

**Figure 3:**
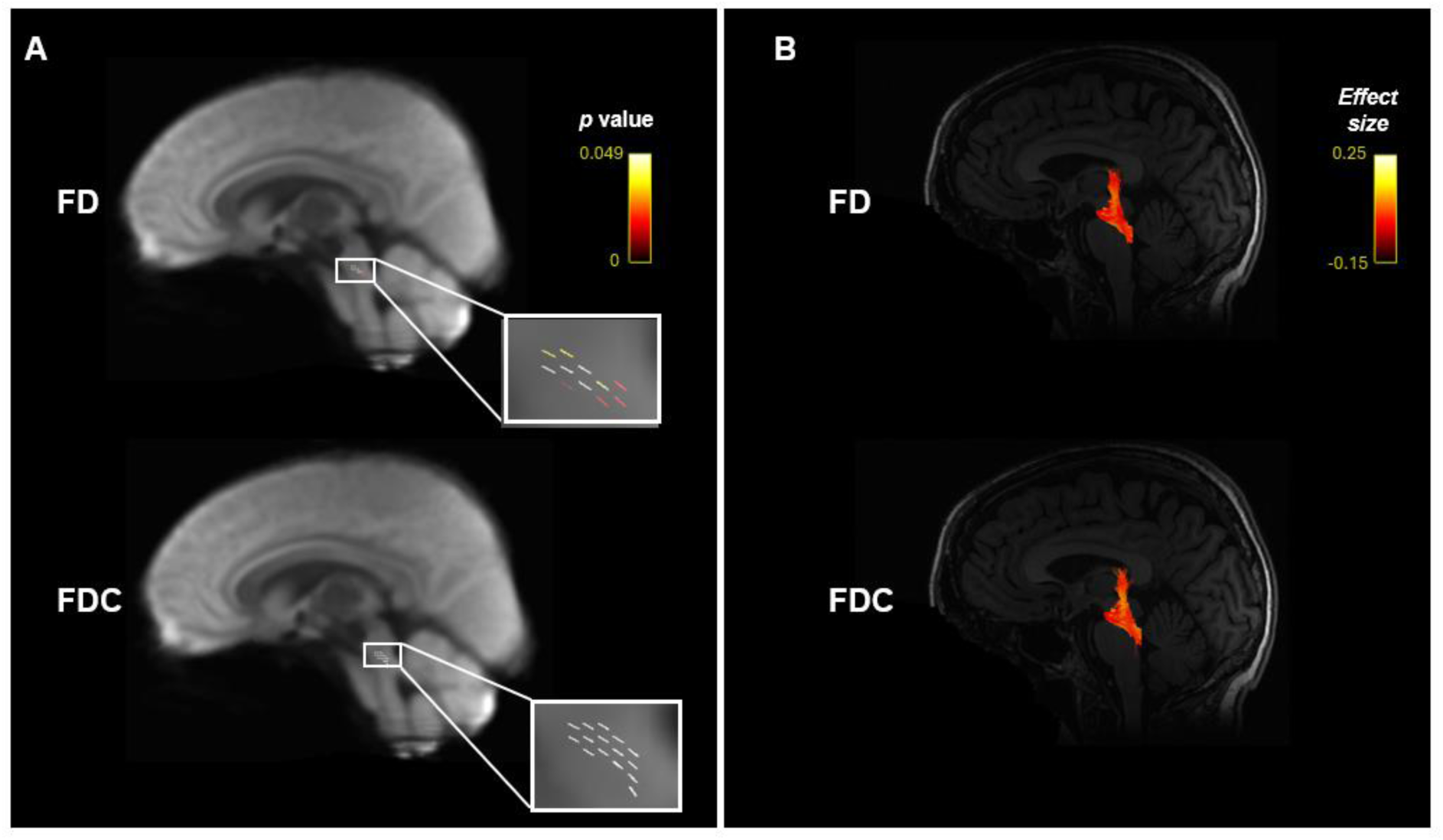
A) Significant Fixel plot Fibre Density (FD), and Fibre Density and Cross-section (FDC) areas overlayed over the population white matter template. B) Streamlines coloured by effect sizes passing through significant Fixel plot FD and FDC areas.

### 3.3 ROI Graph Theory Analyses

Moving from streamlines to white matter networks, between-group differences in ACC were identified for all graph-theory metrics except for global efficiency, which indexes integration over the larger network structure. There were no significant differences (p = 0.115) in global efficiency scores between control (E_glob_=0.063) and R221W carriers (E_glob_=0.068) in higher pain processing network areas. With respect to segregation of white matter networks, R221W carriers had significantly lower local efficiency in the right ACC compared to matched controls. In the control group, the area with highest local efficiency score (reflecting local network segmentation) was the right ACC; in the R221W carrier group it was left somatosensory cortex. Additionally, R221W carriers showed significantly lower betweenness centrality (reflecting the number of shortest paths through a given node) in both right insula and left ACC, compared to controls. Node degree (reflecting the number of edges connecting to a given node) was significantly higher in the right somatosensory cortex and right ACC of R221W carriers when compared to controls. The scores for each graph theory metric are presented in *Table 1*.

**Table 1:**
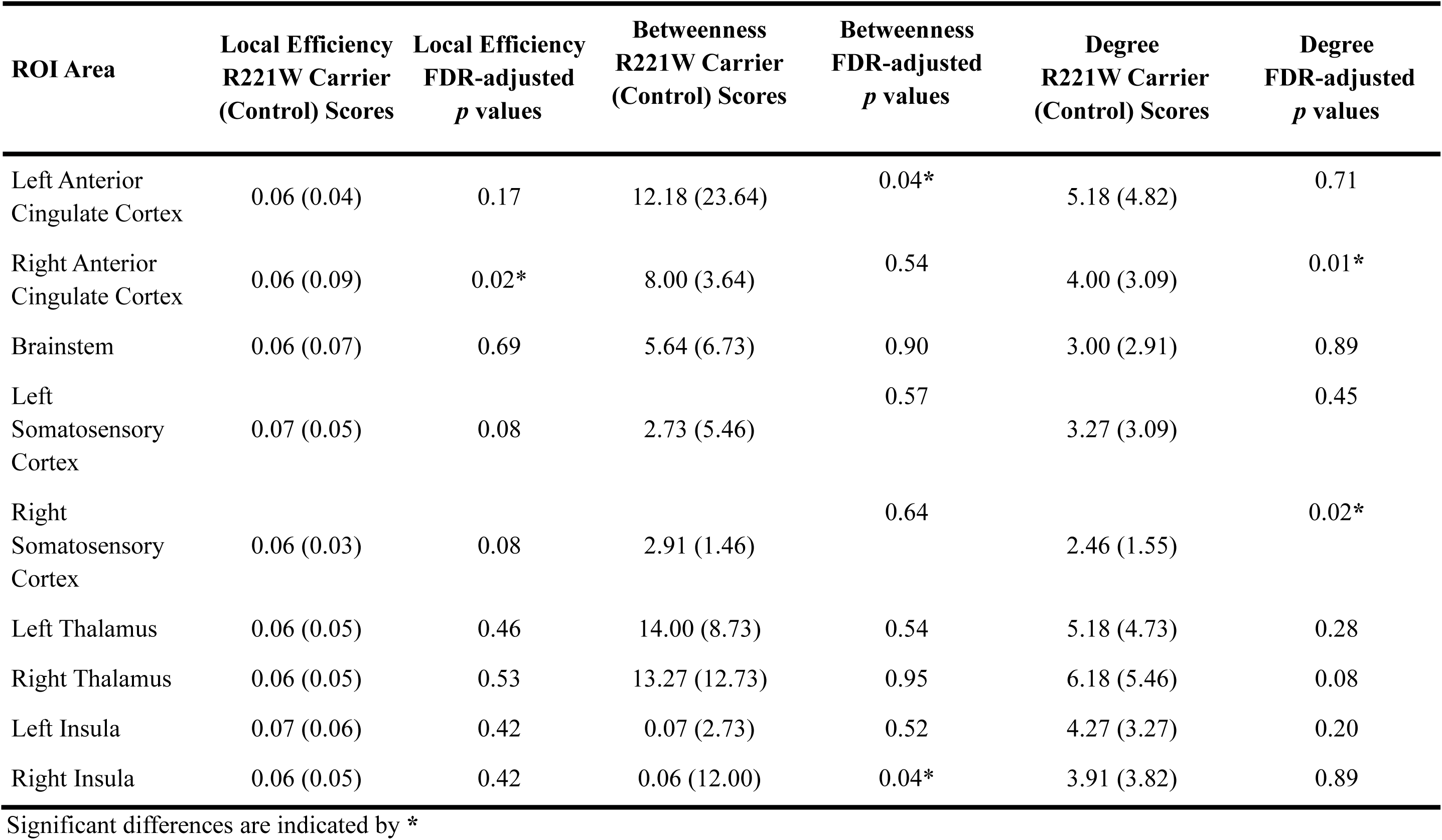
R221 Carrier and Control scores and FDR-adjusted p values of local efficiency, betweenness centrality, and node degree.

| ROI Area | Local Efficiency<br>R221W Carrier<br>(Control) Scores | Local Efficiency<br>FDR-adjusted<br>$p$ values | Betweenness<br>R221W Carrier<br>(Control) Scores | Betweenness<br>FDR-adjusted<br>$p$ values | Degree<br>R221W Carrier<br>(Control) Scores | Degree<br>FDR-adjusted<br>$p$ values |
| --- | --- | --- | --- | --- | --- | --- |
| Left Anterior<br>Cingulate Cortex | 0.06 (0.04) | 0.17 | 12.18 (23.64) | 0.04* | 5.18 (4.82) | 0.71 |
| Right Anterior<br>Cingulate Cortex | 0.06 (0.09) | 0.02* | 8.00 (3.64) | 0.54 | 4.00 (3.09) | 0.01* |
| Brainstem | 0.06 (0.07) | 0.69 | 5.64 (6.73) | 0.90 | 3.00 (2.91) | 0.89 |
| Left<br>Somatosensory<br>Cortex | 0.07 (0.05) | 0.08 | 2.73 (5.46) | 0.57 | 3.27 (3.09) | 0.45 |
| Right<br>Somatosensory<br>Cortex | 0.06 (0.03) | 0.08 | 2.91 (1.46) | 0.64 | 2.46 (1.55) | 0.02* |
| Left Thalamus | 0.06 (0.05) | 0.46 | 14.00 (8.73) | 0.54 | 5.18 (4.73) | 0.28 |
| Right Thalamus | 0.06 (0.05) | 0.53 | 13.27 (12.73) | 0.95 | 6.18 (5.46) | 0.08 |
| Left Insula | 0.07 (0.06) | 0.42 | 0.07 (2.73) | 0.52 | 4.27 (3.27) | 0.20 |
| Right Insula | 0.06 (0.05) | 0.42 | 0.06 (12.00) | 0.04* | 3.91 (3.82) | 0.89 |

Significant differences are indicated by \*

Alongside node differences, edge metrics provided additional information about connectedness between ROI areas. Mean edge betweenness score for controls was *b_e_* = 1.486 (SD = 1.761) and for R221W carriers was *b_e_* = 1.360 (SD = 1.408). Edge betweenness was significantly lower in R221W carriers as compared to controls in left somatosensory cortex connection to the left ACC (p < 0.05), right somatosensory cortex to the right ACC (p < 0.05), and left Insula to the left somatosensory cortex (p < 0.05). Edge betweenness, node degree, betweenness centrality, local efficiency, and edge differences are illustrated in *Figure 4*.

**Figure 4:**
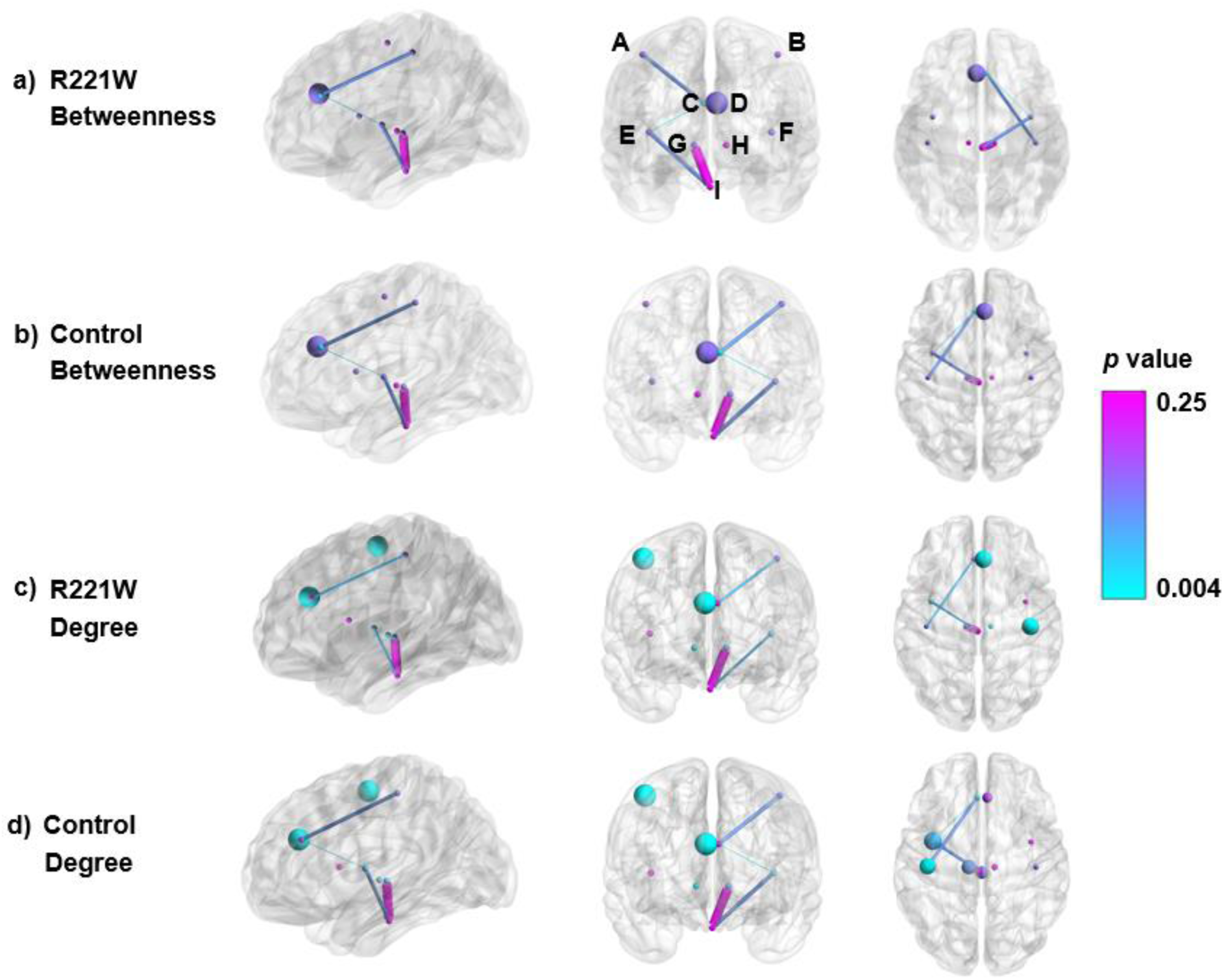
Visualisation of graph theory comparisons. (a) and (b) represent node betweenness of R221W carriers and control groups respectively, with the node size representing effect size and the colour representing their p-value. (c) and (d) represent node degree of R221W carriers and control groups respectively, with the node size representing effect size and the colour representing their p-value. Edge thickness of each group represents edge betweenness score and edge colour represents node efficiency significance scores. Region labels: (A) sensory cortex right, (B) sensory cortex left, (C) ACC right, (D) ACC left, (E) insula right, (F) insula left, (G) thalamus right, (H) thalamus left, (I) brainstem. The figure was made using BrainNet Viewer^65^

### 3.4 ROI NBS Analysis

NBS was performed twice: first to test connectivity of the control group as compared to R221W group, and the opposite comparison for R221W group, all 6 (two sets of 3 t-tests) tests were then corrected for FDR. Similar to graph metrics, both results found significant differences in the thalamus between groups. The R221W group showed weaker connection between the right ACC and right thalamus compared to controls. This difference was only significant at t-test threshold of 3.0. The R221W group did show significantly better connectivity from right thalamus to left ACC and to left insula at all t-test thresholds (*Figure 5*).

**Table 2:**
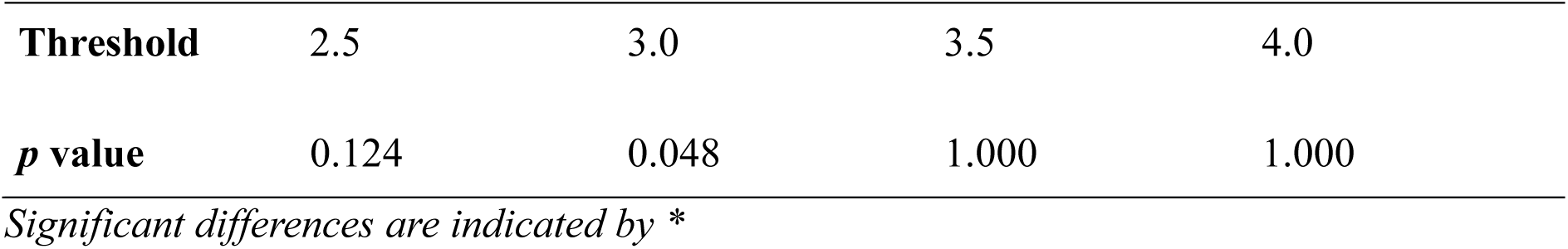
Network Based Statistics comparison of control group against R221W carrier group at different t-test thresholds, multiple comparison corrected.

**Table 3:** Network Based Statistics comparison of R221W carrier group against control group at different t-test thresholds, multiple comparison corrected.

| Threshold | 2.5 | 3.0 | 3.5 | 4.0 |
| --- | --- | --- | --- | --- |
| *p* value | 0.019 | 0.003 | 0.016 | 0.004 |

*Significant differences are indicated by \**

**Figure 5:**
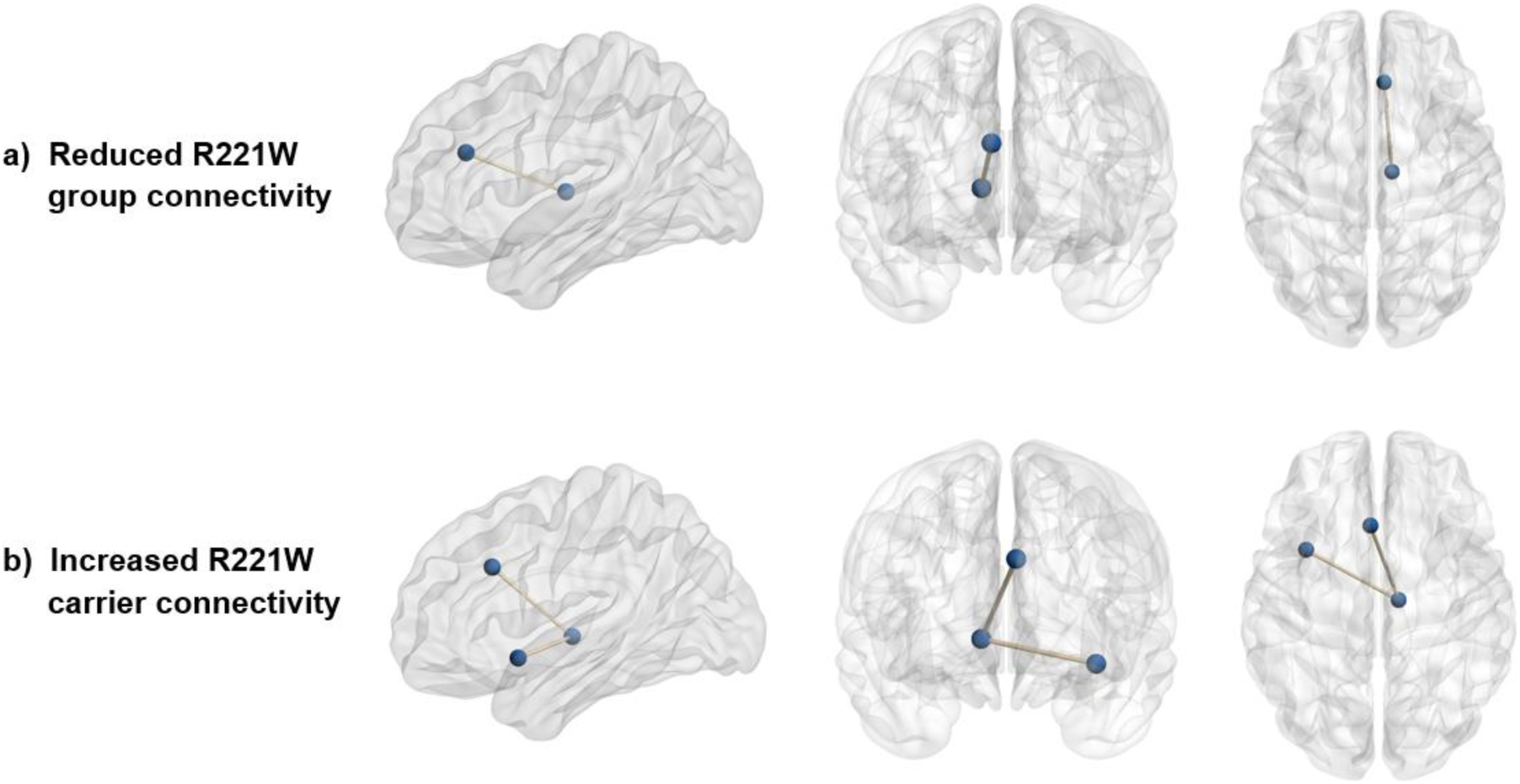
Network Based Statistics significant results represented as nodes and edges

## 4. Discussion

This study provides first evidence that the R221W mutation of the NGF gene is associated with changes to integrity of nerves not only in the peripheral, but also in the central nervous system. Whole brain and ROI Fixel analyses identified fibre density and cross-section reduction in a number of sensorimotor, but primarily motor, white matter tracts passing through the pons and midbrain of R221W carries. The changes being constrained to the brainstem is an important finding as the brainstem is an integral area for processing nociceptive stimulus and affective-motivational pain response^66,67^. These efferent-related changes in brainstem white matter run contrary to our original hypothesis that R221W carriers would display afferent-related spinocortical tract changes, but adds to accumulating evidence that motor^13^, rather than simply somatosensory, changes are integral to the R221W phenotype. Further, although no group differences were found in global network efficiency, which quantifies information integration over the larger white matter network, local network efficiency differences were found in pain-relevant subnetworks including ACC and insula. These local efficiency differences in R221W carriers suggest that inefficient connectivity among these nodes may bias processing to parts of the network more related to somatomotor than affective-motivational aspects of behavioural responses to acute pain.

### 4.1 NGF and the R221W neural phenotype

NGF is primarily constrained to the peripheral nervous system and research on NGF impact on the central nervous system has been limited^68^. Rat cell line models suggest that the missense point mutation involved in the R221W phenotype affects cleavage of pro-NGF in intracellular pathways, limiting the availability of functional mature NGF in the extracellular space^69,70^. The R221W carriers’ reduced unmyelinated C afferent density may result from insufficient trophic support during development, with possible additional effects of altered regulatory signaling in NGF-mediated nociceptive pathways in adulthood. The trophic role of NGF likely does not substantially affect large myelinated Aβ fibre axons, the cell bodies of which are negative for the primary NGF receptor TrkA^71^. However, For brainstem fibre tracts, NGF changes could be coupled with transneural degeneration as a consequence of fewer C- and A-delta fibre inputs^72,73,74^ - second-order neurons that synapse with C-nociceptor fibres in spinal lamina I and extend to parts of the brainstem^75^. Since C-nociceptor fibres are reduced in R221W carriers^13^, congenital disruption of second-order afferent synapses is propagated to downstream connectivity changes in brain networks ^76,77^. These biological changes could influence motor fibres in the brainstem and cortex, and possibly contribute to the reduced behavioural response to pain of R221W^11^.

### 4.2 Somatomotor and affective-motivational network differences in R221W

A majority of the tracts with reduced fibre density and cross section connect the midbrain to the cerebellum and relate to movement coordination: the superior cerebellar peduncle^78^, middle cerebellar peduncle^79^, and the pontine crossing tract^80^. Additionally, effects on corticospinal tract^81^ but not spinothalamic tracts could suggest that the altered behavioural response can be attributed to affected white matter motor efference rather than somatosensory afference. The two other affected tracts, medial lemniscus and inferior cerebellar peduncle, both relate to proprioceptive information relay^82,83^ from the peripheral nervous system to the somatosensory cortex and brainstem, respectively. This information is not nociceptive but could be involved in coordinating motor behaviour in response to pain^84^. These tract differences could relate to the dampened motivated responses of R221W carriers in reaction to painful stimulation^13^. This and previous results are consistent with a transgenic mouse model of the human R221W (or R100W) mutation, which displays behavioural under-reaction and increased response latencies to noxious stimuli, alongside decreased neural activation in motor cortex and striatum^85,86^.

The medial lemniscus projects largely via the ventral posterior inferior thalamus nucleus to primary somatosensory cortex^82^. R221W carriers show reduced fibre density and cross-section here, though their somatosensory cortex and right ACC both had a higher number of connections (node degree) than matched controls. However, the higher number of connections did not translate to higher efficiency, since R221W carriers had significantly lower local efficiency for right ACC, reflecting a lesser degree of segregation among subnetwork connections. The left ACC of R221W carriers also showed fewer connections with immediately neighbouring network nodes, as indicated by reduced node betweenness indexing the number of shortest paths through the node. This raises the possibility that increased node quantity, alongside decreased hallmarks of segregated subnetwork organization in carriers’ somatosensory and cingulate cortices, impinges on network efficiency. Such reduced efficiency in the ACC subnetwork, in conjunction with the highest local efficiency observed in left somatosensory cortex for R221W carriers, could indicate an increased reliance of R221W carriers on the primary somatosensory and motor cortices during acute pain processing.

R221W carriers’ ACC also exhibits reduced integration with wider cortical network regions associated with pain processing, not just to its immediate neighbour areas. That is, R221W carriers appear to have fewer critical links between ACC and the somatosensory cortices in both hemispheres, as indicated by reduced edge betweenness, reflecting a decrease in the overall number of shortest paths through the node relative to controls. Additionally, there was diminished right side lateralised connectivity between the ACC and the thalamus, as R221W carriers had fewer edge connections to the right thalamus. This was also identified in NBS analysis in which the right ACC and right thalamus formed part of a disrupted subnetwork. These laterality differences may reflect a differing degree of bilateral network engagement between carrier and control groups. Such interhemispheric connectivity disruptions may account for pain response latency seen in the anterior-mid cingulate cortex of R221W carriers^13^. In previous research, the BOLD response of R221W carriers’ reported urge to move away from a painful thermal stimulus emerged only after a 2-3 second delay following the stimulus onset, which covaried with MCC activation in controls but not carriers^13^.

### 4.3 Local network reorganisation in the R221W phenotype

The NBS result indicating that R221W carriers have a better-connected subnetwork between the right thalamus, left ACC, and left insula could suggest a compensatory reorganisation not captured in functional MRI research. The increased connectivity between the right thalamus and left ACC could compensate for the reduced connectivity from the left ACC to the somatosensory cortex, as well as providing cross-hemispheric compensation for decreased connectivity between right thalamus and right ACC. This increased connectivity from the right thalamus to the left insula could serve a similar purpose – reorganisation to compensate for the reduced connectivity between left insula to left somatosensory cortex, and cross-hemispheric compensation for the reduced local integration of the right insula. Stronger hemodynamic activation of right AI was proportional to how well R221W carriers differentiated between task-relevant and task-irrelevant painful and nonpainful stimulation^13^. Insula’s functional connectivity with MCC tracked subjective escape urges for painful stimuli in controls but not carriers^12^. To the extent that the insula contributes to task-dependent aspects of pain behaviour, its degree of engagement with cingulate regions in acute pain could bias network dynamics towards more typical behavioural responses.

The stronger connections from right thalamus to the left ACC and left insula could form a subnetwork that works to support the impaired pain motor pathways. The present findings indicate that while the right insula could support adaptive pain behaviour in response to afferent signals from the somatosensory cortex, the left insula could provide compensatory structural integration with the contralateral thalamus. As both the thalamus and insula are associated with motor signal relay and motor control respectively, they could compensate for reduced ACC engagement by motivational aspects of pain in R221W carriers^13^. However, this compensation, alongside increased motor cortex engagement, may contribute to inefficient network processing and attendant latencies in behavioural responses^83^. It is also possible that the interpolation of left insula in this subnetwork renders it less efficient for producing quick and adaptive behaviour. While cross-hemispheric thalamus-insula connectivity cannot fully compensate for impaired ACC integration or impaired motor pathway integrity, these areas could support further downstream signalling among cortical areas responsible for selection, preparation, and execution of responses to noxious stimuli^87,88,89^, albeit in a potentially less efficient or rapid manner.

### 4.4 Limitations and Future Directions

This study has several limitations that warrant consideration. The small sample size, reflective of the rarity of the R221W mutation, inherently limits the generalizability of our findings and may constrain statistical power. To address this, we employed proportional thresholding at 20% of the strongest connections in the structural connectivity matrices during our graph theory analyses. This approach was necessary to ensure comparability across participants by controlling for individual differences in edge density, which is particularly important in studies with small sample sizes^50,51^. However, it is acknowledged that this method may overlook weaker but potentially significant connections, which could be explored in future studies using alternative approaches, such as density-preserving methods.

While the R221W gene mutation group was small, the breadth and impact of these results could have implications for other peripheral neuropathy and pain indifference populations. Further research on other CIP and groups is needed to compare pain insensitivity mechanisms between different congenital conditions, and to compare groups with CIP to acquired peripheral neuropathy on structural brain white matter differences. This comparison is particularly needed as white matter reorganisation has been seen in other congenital sensory loss groups, but it has been found to be significantly different to groups who acquired sensory loss later in life^90,91^.

Although both graph theory and NBS analyses used internal multiple comparisons correction (FDR and permutation-based), the results from each were interpreted independently to minimize the risk of inflated Type I error across analytic frameworks.

For the R221W group, future research will further examine the functional contributions of affective-motivational networks, including the effects of local ACC-AI subnetwork efficiency, with the evidence for a greater reliance on somatomotor-related networks for producing appropriate and timely behavioural responses during acute pain.

### 4.5 Conclusions

This study presents three different white matter analysis frameworks that resulted in nuanced findings of structural disruption and reorganisation in people with CIP. Whole-brain and ROI Fixel-based analyses identified a number of altered R221W carriers’ sensorimotor pathways that could contribute to the reduced behavioural response to pain of R221W carriers. These findings support a previously proposed theory^13^ that CIP in R221W carriers is one of motor under-reactivity and pain indifference, rather than pain insensitivity. Graph theory and NBS identified the cortical somatosensory cortex and motivation-related ACC as having more local connections, but the increased quantity did not translate to increased quality. Somatosensory cortex, ACC, and insula were all affected with respect to local efficiency and global integrity. However, this analysis also identified different structural configurations that could indicate local, global, and cross-hemispheric compensatory reorganisation. Currently, these findings do support the integrative role of the thalamus, ACC, and insula in somatosensory and motor reactivity to nociception^87,88,89^ by showcasing how their disruption can lead to signs of impaired pain processing and reactivity. While there is no previous research of peripheral CIP somatosensory symptoms translating to central nervous system changes, chronic pain neuropathy conditions provide comparable findings of reduced structural integrity in the somatosensory cortex^92,93^ and ACC^94^. These structural findings require further clinical and neuroimaging testing in both CIP and painful peripheral neuropathy populations to understand the full impact of peripheral neuropathy on central nervous system reorganisation. The combined results suggest a complex interplay of disruptions and compensations within pain-processing and motor effector networks, providing support for the theory that congenitally reduced nociceptive fibres can impact the central nervous system development in CIP populations.

## 5. Data and Code Availability

Data is made freely available with CC0 licence on Figshare^95^ and the code is freely accessible on GitHub^96^.

## 6. Author Contributions

Arnas Tamasauskas and Irene Perini contributed equally to this work and are joint first authors. Arnas Tamasauskas: performed data analysis and drafted the manuscript.

Irene Perini: conceived the project, obtained ethical approval, collected data, and contributed to manuscript drafting.

Jan Minde: collected data.

Simon S. Keller: provided guidance on data analysis and critically revised the manuscript.

Nicholas Fallon: provided guidance on data analysis and critically revised the manuscript.

Bernhard Frank: contributed clinical expertise and critically revised the manuscript.

India Morrison: supported data analysis and critically revised the manuscript.

Andrew Marshall: conceived the project, obtained ethical approval, collected data, and supervised the study.

